# Single-stranded DNA library preparation uncovers the origin and diversity of ultrashort cell-free DNA in plasma

**DOI:** 10.1101/035741

**Authors:** Philip Burnham, Min Seong Kim, Sean Agbor-Enoh, Helen Luikart, Hannah A. Valantine, Kiran K. Khush, Iwijn De Vlaminck

## Abstract

Circulating cell-free DNA (cfDNA) is emerging as a powerful monitoring tool in cancer, pregnancy and organ transplantation. Nucleosomal DNA, the predominant form of cfDNA in blood, can be readily adapted for sequencing via ligation of double-stranded DNA (dsDNA) adapters. dsDNA library preparation, however, is insensitive to ultrashort, degraded and single-stranded cfDNA. Drawing inspiration from recent technical advances in ancient genome analyses, we have applied a single-stranded DNA (ssDNA) library preparation method to sequencing of cfDNA in the plasma of lung transplant recipients (40 samples, six patients). We found that the ssDNA library preparation yields a greater portion of sub-100 bp DNA, as well as an increased relative abundance of human mitochondrial cfDNA (10.7x) and microbial cfDNA (71.3x). We report the fragmentation pattern of mitochondrial, nuclear genomic and microbial cfDNA over a broad fragment length range. We furthermore report the first observation of donor-specific mitochondrial cfDNA in the circulation of lung transplant recipients. We found that donor-specific mitochondrial cfDNA molecules are significantly shorter than those specific to the recipient. The higher yield of viral, microbial and fungal sequences that result from the single-stranded ligation approach reduces the cost and increase the sensitivity of cfDNA-based monitoring for infectious complications after transplantation. An ssDNA library preparation method provides a more informative window into understudied forms of cfDNA, including mitochondrial and microbial derived cfDNA and short fragment nuclear genomic cfDNA, while retaining information provided by standard dsDNA library preparation methods.

## Introduction

Cell-free DNA (cfDNA) is quickly finding application as a monitoring tool in pregnancy, cancer and organ transplantation (Fan et al. 2008; Lo et al. 1998a; Diehl et al. 2008; De Vlaminck et al. 2014; Kitzman et al. 2012). cfDNA exists in circulation in many shapes and forms, including fragments of the nuclear genome, the mitochondrial genome and microbial genomes (Jiang et al. 2015). The predominant type of cfDNA is derived from the nuclear genome and has a fragment size centered around 166 bp, approximately the length of a segment of DNA wound around a histone octamer (Mouliere et al. 2011; Quake 2012). These nucleosomal fragments of cfDNA are readily accessible for sequencing using standard library preparation methods that are based on ligation of dsDNA sequencing adapters. The most commonly used implementations of this method rely on multiple bead-based size-selective steps that eliminate unwanted adapter-dimer products. These methods, although relevant to a wide range of applications, are not sensitive to the full diversity of circulating DNA (Mouliere and Rosenfeld 2015), in particular shorter fragments, highly degraded fragments and (partially) single-stranded fragments of DNA in circulation remain undetected (Fig. 1).

**Figure 1.**
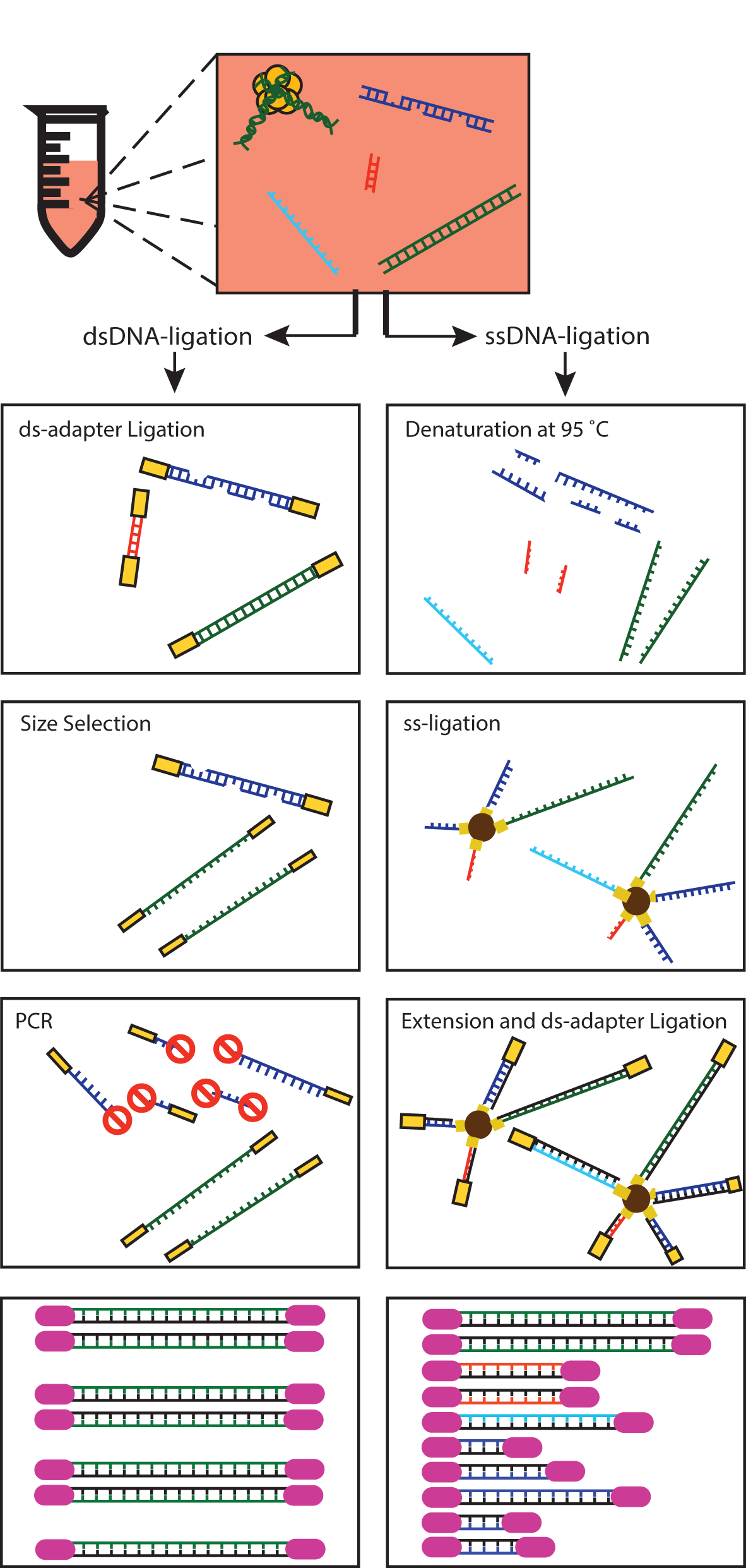
Schematic of sequencing library preparation methods. Schematic illustration of key steps in the dsDNA and ssDNA library preparation protocols used in this work and their sensitivity to different types and forms of circulating cfDNA in plasma. cfDNA in plasma may be single-stranded (light blue), partially single stranded or nicked (dark blue), short (red), or long (green).

An interesting parallel exists with genomic analyses of ancient DNA samples, where the target DNA is usually highly fragmented and present in low amounts (Allentoft et al. 2012). Recently, Gansauge and Meyer introduced a sequencing library preparation method that is based on single stranded ligation and demonstrated the method by sequencing of the genome of an extinct archaic human (Meyer et al. 2012; Gansauge and Meyer 2013). Here, we have applied this protocol to sequencing of cfDNA in plasma, motivated by the hypothesis that an ssDNA library preparation method is, in principle, sensitive to the full diversity of cfDNA in the circulation, including ultrashort dsDNA, ssDNA and dsDNA with nicks in both strands.

We applied the ssDNA library preparation to the analysis of clinical samples collected from organ transplant recipients and compared data of fragment types, lengths and abundance to results from dsDNA ligation assays performed on the same plasma DNA extracts (De Vlaminck et al. 2013, 2015). Organ transplantation is an interesting model system for this work, given that both graft- and patient-derived DNA is present in plasma, providing a window into the tissue specificity of the properties of cfDNA. Furthermore, transplant recipients are subject to immunosuppressive therapies that reduce the risk of rejection, but increase their susceptibility to opportunistic infections. Analyses of microbial cfDNA in plasma are therefore particularly relevant in this context and can be analyzed in conjunction with clinical data. We note that Karlsson et al. recently applied an ssDNA ligation protocol to the amplification-free sequencing of cfDNA (Karlsson et al. 2015). These authors, however, did not perform an analysis of fragment lengths and did not examine the presence of mitochondrial and microbial DNA in plasma.

## RESULTS

Forty samples of cfDNA extracted from plasma of six double-lung transplant recipients (De Vlaminck et al. 2015) were analyzed in this study. Sequencing libraries were prepared using a single-stranded DNA library preparation protocol and sequenced (5.7 ± 1.4 million paired-end reads per sample). Results were compared against sequence data obtained for the same samples following double-stranded DNA library preparation where available (36 matched samples, 18.8 ± 9.1 million paired-end reads per sample). The key distinguishing features of the library preparation protocols are schematically represented in Figure 1.

We analyzed data using two distinct workflows. The first eliminates human-aligned sequences and assigns non-human fragments to annotated microbial references using BLAST (De Vlaminck et al. 2013). The second workflow aligns raw reads to the human genome [GenBank:GCA_000001305.2] extended with an edited human mitochondrial reference genome [GenBank:NC_012920] (see SI).

### Size distribution and abundance of mitochondrial DNA in plasma measured by digital PCR

A retrospective analysis of sequencing data of cfDNA in plasma of lung transplant recipients available from a previous study (De Vlaminck et al. 2015) revealed a fractional abundance of mitochondrial cfDNA of 2×10^−3^ %, which is in line with a recent observation, but is low considering that there are 50-4,000 mitochondria per cell (Miller et al. 2003). We used digital PCR (dPCR) assays with varying amplicon length (49-304 bp) to characterize the abundance of mitochondrial cfDNA prior to library preparation and compared this to the size distribution of nuclear genomic cfDNA (Fig. 2A). The experimental design with variable amplicon lengths provided information about the underlying fragment length distribution (Mouliere et al. 2011): cfDNA is randomly fragmented in plasma - the genomic abundance, as measured by PCR, is therefore expected to decrease monotonically with amplicon length, with a gradient that is a function of the underlying fragment length distribution. These experiments revealed that mitochondrial DNA is more fragmented than nuclear genomic DNA (mean fragment length < 100bp), but present at much greater abundance in plasma (56-fold greater representation, genome equivalents). The consequence of the short fragment size of mtDNA is that commonly used dsDNA library preparation protocols, which require multiple bead-based size-selective steps that eliminate unwanted adapter-dimer products, are relatively insensitive to mitochondrial sequences.

**Figure 2.**
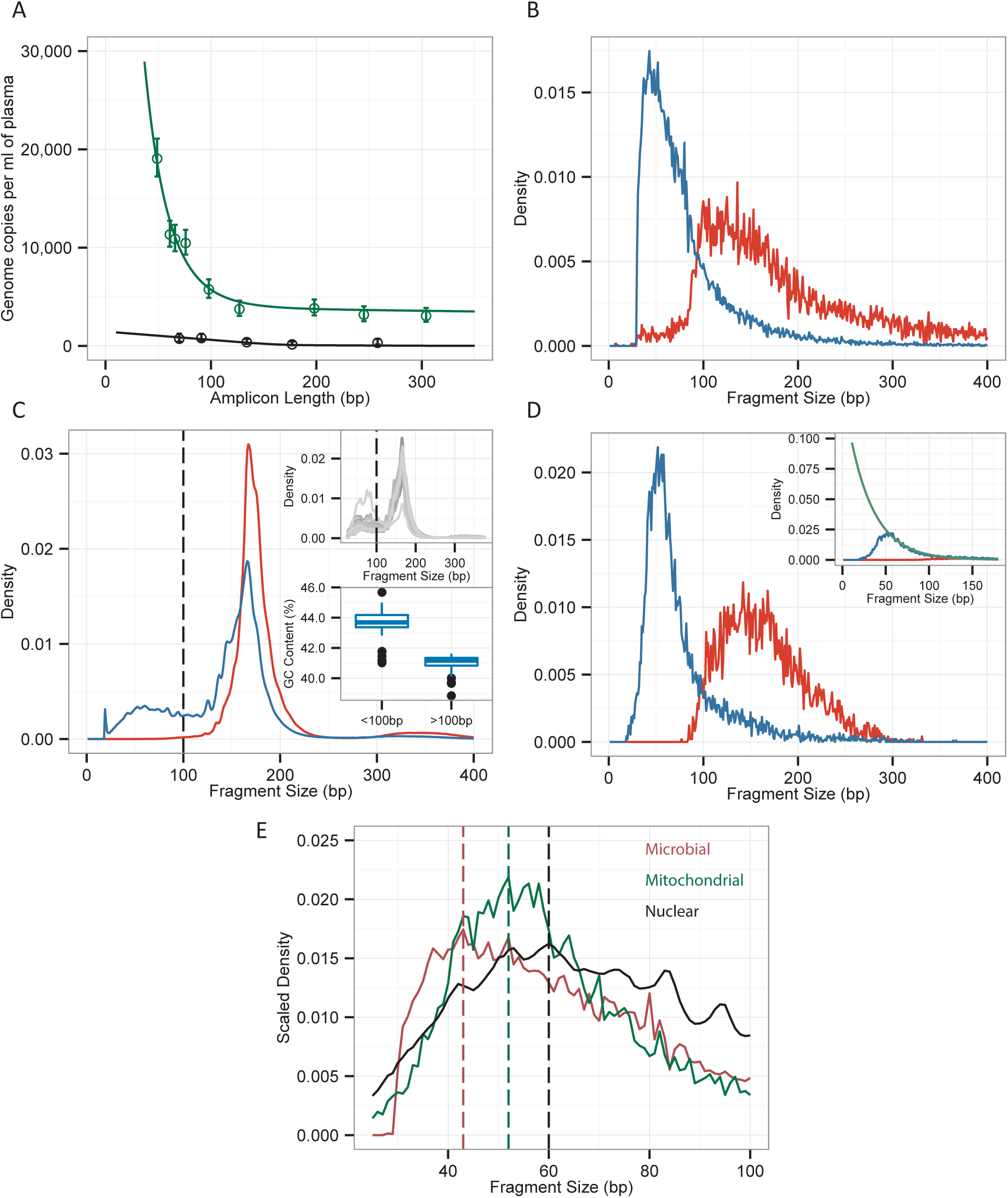
cfDNA fragment length distributions. **(A)** Abundance of mitochondrial (green) and nuclear genomic cfDNA (black) measured by digital PCR assays with different amplicon lengths. Solid lines are model fits (see SI). **(B-D)** Fragment length histograms measured via sequencing for microbial **(B)**, nuclear **(C)** and mitochondrial genomic **(D)** cfDNA following ssDNA (blue) and dsDNA (red) sequencing library preparation. Inset of **(C)** Indicates the sample-to-sample variability of fragment lengths measured, as well as the difference in GC content for short (< 100 bp) and long (> 100 bp) regions marked with the dashed line. **(E)** Density (scaled for clarity) of short length (segment lengths < 100 bp) mitochondrial, microbial and nuclear genomic cfDNA measured by ssDNA library preparation. Vertical lines highlight most prevalent fragment lengths.

### Library preparation via single-stranded ligation

We implemented a single-stranded DNA library preparation protocol first described by Gansauge and Meyer (Gansauge and Meyer 2013) that does not require size-selective steps that eliminate shorter fragments (Fig. 1 and Methods). A positive control (1 μL of 500 μM, synthetic ssDNA) and a negative control were included with each batch of samples. The efficiency of ligation of cfDNA fragments to biotinylated probes and ligation of double stranded adapters to primer-extended products was measured using quantitative PCR (as in (Gansauge and Meyer 2013)). On average 0.83×10^9^ unique ssDNA molecules (0.016 to 8.2 ×10^9^) were ligated and underwent PCR amplification (8 to 15 cycles) prior to sequencing. The concentrations of sequencing libraries were 3.45 ng/μL ± 1.72 ng/μL. Using the positive control as a measure, we determined that, on average, 22.2% of cfDNA molecules bind to biotinylated adapters, and 15.1% of those that are biotinylated survive the rest of the procedure, which includes double-stranded adapter ligation and extension; as a result, 3.35% were PCR amplified and sampled for sequencing. By adjusting the extension sequence primer with a 4-N overhang on the 5’ end (as in (Karlsson et al. 2015)), we limited the occurrence of adapter dimers to, on average, one in 1,700 sequences.

### DNA Fragmentation profiles

We used paired-end sequencing to determine the fragment lengths of nuclear, mitochondrial and microbial cfDNA. Briefly, paired-end sequences were aligned (BWA-mem, (Li and Durbin 2009)) and the insert lengths were deduced from the coordinates of the bases at the outermost ends of each sequence pair. For microbial sequences, fragments assigned to microbial genomes using BLAST (Altschul et al. 1990), were retrieved and realigned using BWA-mem to determine the lengths of the library inserts.

Figures 2B-2D show a direct comparison of the fragment length profiles measured for the two library preparation methods. We find that DNA shorter than 100 bp becomes much more accessible for sequencing following ssDNA library preparation. While dsDNA library preparation results in detection of only a few molecules of mitochondrial and microbial cfDNA with length shorter than 1000bp, the use of ssDNA library preparation revealed an abundance of such molecules with lengths between 50 and 100 bp. The lower limit of detection for ssDNA libraries was ~40 bp for all subclasses of cfDNA (mitochondrial, microbial, nuclear genomic cfDNA), pointing to a limit set by the DNA isolation method (Fig. 2E).

The peak in the length profile at ~160-166 bp for cfDNA fragments assigned to the nuclear genome (Fig. 2C) is likely a consequence of the protection of these molecules from degradation by nucleases in the blood through tight association with histones. This property has been reported in many previous studies and is observed for both the ssDNA and dsDNA library preparation protocols (Schwarzenbach et al. 2011). A second peak at shorter lengths (< 100 bp) is unique to the libraries prepared by single-stranded ligation (De Vlaminck et al. 2013). The relative proportion of nuclear genomic DNA shorter than 100 bp made up a substantial proportion of nuclear cfDNA (20.54% ± 11.51%). We partitioned non-nucleosomal DNA into two groups, those with length under and over 100 bp, to examine distinguishing features between the two groups. The GC content between the groups differed significantly (Fig. 2C, inset); the GC content of the super-100 bp group was 40.9%, while that of the sub-100 bp group was 43.5%. We hypothesized that the sub-100 bp nuclear cfDNA may be derived from coding regions of genes, which are known to be GC-rich (Vinogradov 2003; Ivanov et al. 2015). These observations indicate that a considerable amount of nuclear genomic cfDNA is not nucleosome protected and, thus, subject to degradation by nucleases in the blood.

Previous reports suggest that fetal and tumor derived cfDNA are shorter than cfDNA derived from maternal and normal tissue - the sensitivity of the ssDNA library preparation protocol to molecules over a wider length range is therefore a feature that will be useful for applications in prenatal testing and tumor monitoring.

### Improved recovery of mitochondrial and microbial cfDNA

We next examined the coverage of mitochondrial and microbial genomes relative to the nuclear genome for the dsDNA and ssDNA library preparations. We found that the ssDNA library preparation gives rise to an increase in the relative number of mitochondrial sequences in the datasets (10.7x) and an increase in the relative coverage of the mitochondrial genome (7.22x). This observation is consistent with the greater sensitivity of the ssDNA library preparation to short fragment DNA described above. The mitochondrial-assigned cell-free DNA (mt-cfDNA) distributions for the ssDNA and dsDNA library preparation methods are highly similar for fragment lengths longer than 130 bp. The fragmentation profile of mitochondrial cfDNA measured by ssDNA ligation is well described with a single exponent (inset Fig. 2D, best fit *a*\*exp(*− c x*); *a* = 1207, *c* = 28.72 bp−^1^,).

To study the efficiency of recovery of microbial derived cfDNA, we estimated the genome coverage of microbes detected across all samples relative to the coverage of the human genome. We next compared the relative genomic coverage of strains detected by both methods in matched samples (n=36) (Fig. 3A); for example E. coli in lung transplant patient L77 on day 3 was treated as a separate event from E. coli in the same patient on day 2. We examined over 1,100 direct comparisons and found a good correlation in the relative genomic abundance as measured by the two distinct library preparations (Fig. 3A). Importantly, library preparation by ssDNA ligation gave rise to a 71-fold increase in the relative genomic coverage of microbial species (74-fold for bacteria, which made up 89% of the sampled species comparisons; see SI, Table 1, Fig. 3B). Consistent with the greater recovery efficiency of the ssDNA protocol, we find that most of the species detected in the dsDNA library preparation assays were also detected following ssDNA library preparation (95% species recovery, 934/984, Fig. 3C). 55% of all species detected were uniquely observed in the ssDNA library preparation assays.

**Figure 3.**
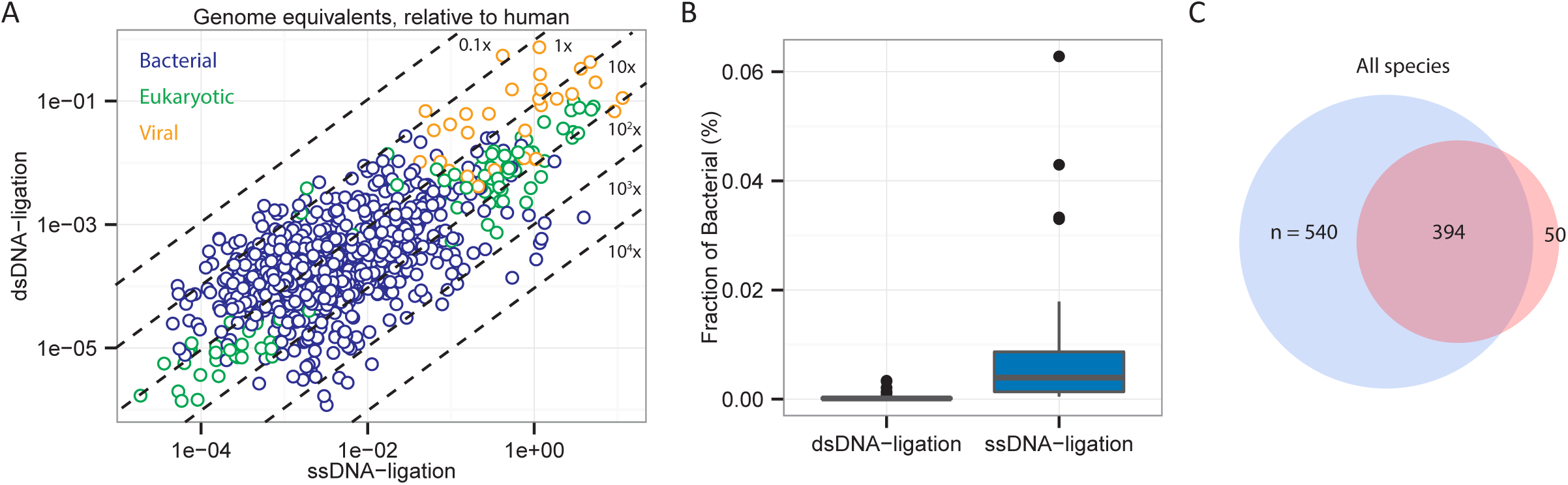
ssDNA library preparation yields greater fraction of non-human cfDNA. (A) Comparison of the coverage of microbial genomes relative to the human genome for ssDNA and dsDNA library preparation. Data points are colored by domain of life. (B) Yield of bacterial sequences for ssDNA library preparation relative to dsDNA library preparation (74-fold mean increase). (C) Venn diagram representation of the number of species uniquely detected following ssDNA library preparation in blue (540/984, 54.9%), species uniquely detected following dsDNA library preparation in red (50/984, 5.1%), and species detected following both protocols (394/984, 40.0%).

The greater efficiency in recovery of microbial cfDNA is in line with the greater sensitivity of the ssDNA library preparation protocol to short fragment DNA. This feature offers the potential to profile the bacterial and viral components of microbiome in plasma more effectively and to perform infectious disease diagnostics based on sequencing of cfDNA with increased precision and at lower cost. Given the vastly different fragmentation profiles of nuclear genomic and microbial cfDNA, it will be possible to further improve the relative abundance of microbial sequences in datasets through size selection strategies.

### Donor-specific cell-free DNA

Transplant donor-specific cfDNA is present in the circulation of organ transplant recipients (Lo et al. 1998b) and recent studies have shown that the proportion of donor-specific cfDNA (cfdDNA) is predictive of acute rejection in heart and lung transplantation (Beck et al. 2013; De Vlaminck et al. 2014, 2015). We compared the fractional abundance of cfdDNA in the lung transplant samples measured following dsDNA and ssDNA library preparation (36 matched samples, six patients, Fig. 4A). We found an excellent agreement between matched measurements (Pearson, c=0.98, p<<1×10^−5^). Here, sequences were assigned to the donor and recipient based on genotype information (single-nucleotide polymorphisms, SNPs) obtained from pre-transplant whole blood samples (De Vlaminck et al. 2015). Interestingly, we compared the fraction of short fragment nuclear genomic cfDNA (segment lengths shorter than 100 bp) to the fraction cfdDNA and found they are correlated (Spearman, ρ=0.692, p<1×10^−5^).

**Figure 4.**
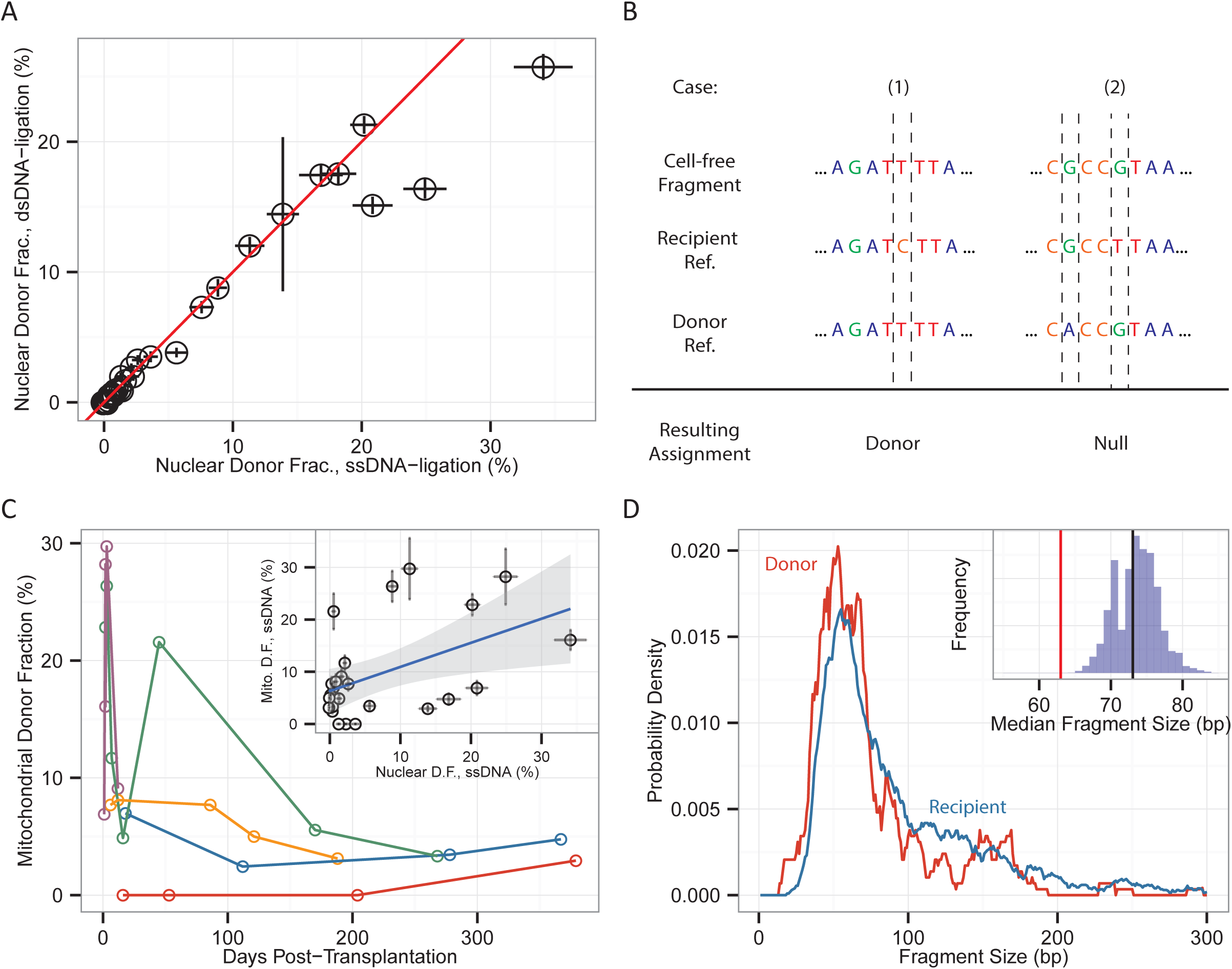
Quantifying donor-specific nuclear genomic and mitochondrial cfDNA. **(A)** Comparison of the fraction of donor-specific nuclear genomic DNA measured after dsDNA and ssDNA library preparation. **(B)** Schematic representation of analysis workflow used to discriminate donor and recipient specific mt-cfDNA. Examples of an ambiguous assignment and a fragment assigned to the donor are shown. **(C)** Fraction of donor-specific mt-cfDNA as function of time post-transplant for five double lung transplant patients (25 samples, having excluded samples with fewer than 20 informative mitochondrial fragments); the inset compares the fraction of donor-specific mitochondrial and nuclear genomic DNA for the same samples (Pearson, c = 0.463, p = 0.0196). **(D)** Smoothed (distribution five nearest-neighbors, running mean) of donor mt-cfDNA is compared to the smoothed distribution of recipient mt-cfDNA. Inset: median fragment size for the donor mt-cfDNA compared to the fragment size of 10,000 subsets sampled from the recipient mt-cfDNA length set.

Because of the high copy number of mtDNA in cells and the relatively high genetic diversity between two unrelated individuals (Aquadro and Greenberg 1983; Li and Sadler 1991), mtDNA is often used in forensic analyses (Linch et al. 2001) and in studies of population genetics (Avise et al. 1987). The same attributes make mitochondrial DNA a promising candidate marker of post-transplant graft injury. We therefore asked whether donor-specific mitochondrial DNA can be detected in the plasma of transplant recipients. We built mitochondrial reference sequences to assign mitochondrial cfDNA to the transplant donor or recipient. To this end, DNA was extracted from whole blood samples collected from the donor and the recipient prior to the transplant procedure. Mitochondrial DNA was selectively amplified and sequenced. One million sequences led to a per-base coverage greater than 100-fold (genome size 16.5 kb), sufficient to determine subject-specific mitochondrial variants. Based on the reference sequences, we compiled lists of SNPs that are unique to either the donor or recipient (Fig. 4B). On average, 152 informative SNPs were found per donor-recipient pair, roughly leading to a SNP every 114 bp. For samples prepared via ssDNA ligation, 8.7% ± 3.4% of the mitochondrial sequences were informative, and 9.5% of the informative SNPs were assigned to the donor.

To the best of our knowledge, this is the first direct observation of graft derived mitochondrial DNA in the circulation of transplant recipients. We computed the fractional abundance of donor-specific mt-cfDNA as the number of donor-specific mt-cfDNA molecules divided by the total number of informative mt-cfDNA molecules. We studied the variability and time dependence of the levels of donor-specific mt-cfDNA. The fraction of donor-derived mt-cfDNA was elevated during the first month post-transplant (Fig. 4C), in keeping with previous observations of elevated levels of cfdDNA in heart and lung transplant recipients during the first few weeks post-transplant, in the absence of acute rejection (De Vlaminck et al. 2014, 2015). The fraction of donor-specific mt-DNA was only modestly correlated (Pearson correlation 0.480, p=0.0152) with the fraction of nuclear genomic DNA. Samples for which there were less than 20 informative mitochondrial fragments were removed (11/36 samples) though the correlation of metrics was relatively stable with decreasing sample size (see Fig. S1). Deeper sequencing and an analysis of a greater set of samples will be needed to investigate the relationship between acute rejection and the release of mitochondrial DNA from the graft.

Previous studies have found differences in fragment lengths for fetal and maternal cfDNA (Tsui et al. 2012), tumor and somatic cfDNA (Mouliere and Rosenfeld 2015) and hematopoietic and non-hematopoietic cfDNA (Jiang et al. 2015). Here, we compared the length of mitochondrial fragments derived from the graft (n = 265) to those specific to the recipient (n = 1855, 40 samples). We generated 10,000 random subsamples of the total collection of recipient-specific fragments (subsampled to the total number of donor fragments detected, n = 265). We next computed the median lengths for the random subsamples and compared to the median length of donor fragments (inset Fig. 4D). We found that donor sequences were slightly shorter (−9 ± 3 bp) than recipient-specific mitochondrial sequences (Fig. S2). This shortening in fragment length may be indicative of differences in the mechanisms of release, or differences in processes of degradation, of donor and recipient mt-cfDNA.

## DISCUSSION

In this work, we have demonstrated that an ssDNA sequencing library preparation method is sensitive to cfDNA of a broad range of types and lengths. Few studies have focused on ultra-short cfDNA (with lengths shorter than 100 bp) or cfDNA that is not derived from the nuclear genome, including mitochondrial and microbial derived cfDNA. The present work indicates that these relatively overlooked forms of cfDNA provide a unique window into physiology.

We applied the ssDNA library preparation method to the analysis of cfDNA in the plasma of lung transplant recipients. We report the first observation of graft-derived mitochondrial DNA in the plasma of these organ transplant recipients. Donor-derived mitochondrial cfDNA has not been investigated as a marker of acute rejection in solid-organ transplantation, but offers several advantages: (1) the mitochondrial genome is small and relatively straightforward to characterize via sequencing, (2) the mitochondrial genome contains a great number of variants that enables differentiation of donor and recipient sequences, and (3) with thousands of copies of mitochondrial DNA present in every cell, mitochondrial cfDNA is abundant in plasma. Mitochondrial DNA has conserved similarities to bacterial DNA and contains inflammatogenic unmethylated CpG motifs (Zhang et al. 2010). It is therefore not surprising that mitochondrial DNA was identified as a powerful damage associated molecular pattern - an endogenous molecule that can activate innate immunity when released during cellular injury (Oka et al. 2012). It is conceivable that the release of mitochondrial DNA that accompanies graft injury promotes many of the harmful immunologic responses observed in solid-organ transplantation. The results presented here provide the first window into this relationship.

Microbial DNA that is the product of microbial degradation across the body or of microbes that infect the blood is found circulating in plasma (Dinakaran et al. 2014). We found that the ssDNA library preparation is more effective at recovering bacterial and viral cfDNA, as compared to a dsDNA library preparation method. We furthermore found that the fragmentation length profiles of microbial and mitochondrial DNA in plasma are highly similar, indicating that they are exposed to similar degradation processes. These observations enable measurement of the bacterial and viral microbiome in plasma with greater sensitivity and at a reduced cost.

Previous studies of the molecular size of nuclear genomic cfDNA have provided insight into the origin and nature of these molecules (Van Der Vaart and Pretorius 2008). Many studies have noted that the predominant fragment size of cfDNA is consistent with the size of DNA wrapped around a single histone octamer. Distinct length profiles are observed for cfDNA depending on their cellular origin with hematopoietically derived DNA being longer than nonhematopoietically (Tsui et al. 2012). Here, we found that the sequencing library preparation method can have a huge effect on length profile measurements. We report the fragmentation profile of nuclear genomic cfDNA in plasma over a broad range of lengths, and we conclude that a considerable fraction of nuclear genomic cfDNA is non-nucleosomal and subject to degradation by nucleases, in much the same way that we described for mitochondrial and nuclear genomic cfDNA.

In its current implementation, the ssDNA ligation requires more hands-on time compared to standard protocol (~15 hours versus ~6 hours, for 12 samples) at a similar cost per sample ($35-$40). This work focused on the cfDNA in plasma, but the methods described herein will further be relevant for genomic measurements of cfDNA in urine (Tsui et al. 2012). The widespread interest in circulating cfDNA as a marker of disease, warrants further investigation into the properties, types and origins of cfDNA and motivates further advances in genomic measurement techniques.

## METHODS

**Study Design and sample collection**. We performed additional analyses and experiments on samples collected from six double lung transplant recipients in the scope of a previous study (De Vlaminck et al. 2013, 2015). Twelve whole blood samples that were collected prior to transplantation and 40 plasma DNA samples collected longitudinally after the transplant were analyzed. Here, plasma was extracted via centrifugation and cfDNA was purified from plasma (QIAamp Circulating Nucleic Acid kit) as previously described (De Vlaminck et al. 2014).

**Sample Processing**. Pre-transplant whole blood samples were processed to create consensus mitochondrial sequences for the six transplant donors and six transplant recipients. DNA was extracted from whole blood using the Qiagen DNeasy Blood & Tissue kit (Cat. No. 69581). Mitochondrial DNA was selectively amplified (Qiagen REPLI-g Mitochondrial DNA kit, Cat. No. 69581), and sheared to 300 bp (Covaris). Libraries were prepared for sequencing using the NEBNext Ultra library preparation kit (New England Biolabs, Cat. No. E6040L). The libraries were characterized using the Agilent BioAnalyzer for fragment lengths, quantified by quantitative PCR and sequenced (2×250bp, Illumina MiSeq).

**Plasma DNA samples and processing**. Plasma was extracted via centrifugation and cfDNA was purified from plasma as described previously (QIAamp Circulating Nucleic Acid kit). ssDNA sequencing libraries were prepared from cfDNA purified from plasma following the protocol described by Gansauge and Meyer (Gansauge and Meyer 2013) with the following exceptions: (1) uracil excision steps using endonuclease VIII were not performed, (2) the amount of CircLigase II enzyme in the protocol was reduced from 4 μL to 0.8 μL and amounts of MnCl_2_ and CircLigase II buffer were halved, (3) extension primer CL9, had an addition N*N*N*N overhang on the 5’ end (described in (Karlsson et al. 2015)) to prevent formation of adapter-dimers, (4) all oligonucleotides were purchased through IDT and prepared with standard desalting, not HPLC, and (5) a thermal shaker was not used; rather, heat blocks containing samples were regularly vortexed to keep magnetic beads suspended. Following completion of libraries, samples were sequenced (2×75 bp, Illumina MiSeq or HiSeq).

**Fragment length measurements by digital PCR**. We determined the abundance of mitochondrial and nuclear genomic cfDNA fragments of various sizes via digital PCR (QuantStudio). Whole blood was obtained from a dog and cfDNA was isolated. A panel of amplicons for various sizes from 49 to 304 bp was created using IDT Custom Oligonucleotide Synthesis (see Si for primer design). Forward and reverse primers (each 10μM, 0.3 μL) and cfDNA (2 μL) were mixed with 3.4 μL H_2_O, 7.5 μL QuantStudio 3D Digital PCR Master Mix (2X) (ThermoFisher Scientific, Cat. No. 4485718), and 1.5 μL of SYBR Green PCR Master Mix (ThermoFisher Scientific, Cat. No. 4309155). We performed PCR under the following conditions: (1) 96 °C for 10 min, (2) 55 °C for 2 min, (3) 98 °C for 30 sec, (4) repetition of (2)-(3) for 39 cycles, (5) 55 °C for 2 min, (6) 10 °C hold.

**Analysis workflow to build mitochondrial reference sequences**. Fastq files were trimmed (Trimmomatic, LEADING:25 TRAILING:25 SLIDINGWINDOW:4:30 MINLEN:15,(Bolger et al. 2014)) and aligned against the human reference genome [GenBank:GCA_000001305.2] using BWA-mem (Li and Durbin 2009). Sequences that mapped to the mitochondrial reference sequence (edited from [GenBank:NC_012920], described in Supplementary Information and Fig. S3) were extracted. A BCF file of SNPs was created and a FASTA consensus sequence was determined. A list of informative SNPs was created through comparison of the donor and recipient consensus sequences.

**cfDNA analysis workflow**. Raw sequencing datasets were trimmed (Trimmomatic, LEADING:20 TRAILING:20 SLIDINGWINDOW:4:20 MINLEN:25, (Bolger et al. 2014)), and low quality reads were filtered (FASTX toolkit, -q 21 -p 50, (Gordon and Hannon 2010)) and aligned (BWA-mem, (Li and Durbin 2009)) to the human reference genome [GenBank:GCA_000001305.2], with changes made to the mitochondrial genome described in the SI section. Sequences that mapped to the mitochondrial reference [GenBank:NC_012920] were collected and SNPs were listed using SAMtools (Li et al. 2009). Sequences with SNPs from the standard reference were assigned to the donor or recipient through comparison to the list of informative SNPs compiled for the donor-recipient pair. The analysis workflow used to quantify the fraction of donor-derived nuclear genomic cfDNA has been previously described in detail (De Vlaminck et al. 2014; Snyder et al. 2011). The analysis workflow used to quantify non-human derived sequences has also been described (De Vlaminck et al. 2013).

**Method comparison**. Samples were compared against the same plasma samples that were prepared via a dsDNA library preparation and published computational workflow (De Vlaminck et al. 2013). Raw sequencing data of identical samples, experimentally processed through both standard and ssDNA-ligation methods, was analyzed using the same parameters.

**DATA ACCESS**. Sequencing data are available in the Sequence Read Archive, BioProject 306662 (Public on 1/11/2015,) including BioSamples SAMN04359448 though SAMN04359535.

## ACKNOWLEDGMENTS

We thank Fanny Chen and Peter Schweitzer for assistance with library preparation and sequencing assays. We thank Erin Berthelsen, Robert Goggs, Elizabeth Wilcox and Rory James Todhunter (Cornell University College of Veterinary Medicine), for providing animal blood samples for assay testing and development. This work is supported by a Cornell Startup grant and the Noyce Foundation.

## CONTRIBUTIONS

PB conceived of study, performed experiments, analyzed data, and prepared manuscript. MSK analyzed data. HAV and SAE provided insights and discussion. HL, HAV and KKK provided human samples and organized patient recruitment. IDV conceived of study, analyzed data, and prepared manuscript. All authors read and approved the final manuscript.

## DISCLOSURE DECLARATION

Cornell University has applied for a patent relating to methods described in this study.

**Description of additional file**. Additional data file containing supplementary information and figures are provided in “Burnham_ShortDNA_Supplement.pdf” these include: a description of changes made to the reference mitochondrial genome, a calculation for the genomic abundance from digital PCR data, a table for the relative abundance increase in microbial strains (Tab. S1), a list of oligonucleotide sequences used in dPCR (Tab. S2), the effects of reducing sample size on mt-cfDNA donor fraction (Fig. S1), and a comparison of the coverage of the donor and recipient mt-cfDNA fragments across the mitochondrial genome (Fig. S2), and an analysis of SNP and sample mt-cfDNA donor fractions (Fig. S3).

